# Quantifying Epithelial Plasticity as a Platform to Reverse Epithelial Injury

**DOI:** 10.1101/2020.01.14.906008

**Authors:** Kristine Nishida, Baishakhi Ghosh, Lakshmana Chandrala, Saborny Mahmud, Si Chen, Atulya Aman Khosla, Joseph Katz, Venkataramana K. Sidhaye

**Author notes:** Department of Pulmonary and Critical Care Medicine, Shanghai East Hospital, Tongji University, Shanghai, China (Si Chen). K. Nishida and B. Ghosh contributed equally to this work.

## Abstract

Epithelial surfaces lining the lung serve as the primary environmental gaseous interface, and are subject to common life-limiting diseases, including COPD (Chronic Obstructive Pulmonary Disease). Despite the critical role of epithelial cells in pulmonary health and disease, quantitative models are lacking but are required given the large patient to patient variability to characterize the epithelial plasticity that follows injury. We have identified a series of assessments to quantitatively identify the changes that occur in the epithelium and to identify targets that reverse injury. The injured epithelium has decreased ciliary function and monolayer height, which in the case of cells derived from COPD patients results in an overall disorganization of structure. Injury causes the cells to shift to an unjammed state, with corresponding increases in the velocity correlation length implicating cell shape and stiffness as fundamental to the injury response. Specific inhibitors of actin polymerization (LatA), of MAPK/ERK kinase (U0126) and Nrf-2 pathway activation (CDDO-Me) push the epithelium back towards a jammed state with decreased cell movement and correlation length, as well as improve barrier function and CBF. These studies attest to cell intrinsic properties that allow for a transition to an unjammed state, and that quantitative phenotypic analysis can identify potential specific pharmacologic targets in a given patient and provide insight into basic mechanisms of cellular damage.

**One Sentence Summary:** Environmental toxins undermine tissue integrity by manipulating transitions from jammed to unjammed states, thereby mimicking or inducing disease.

## Introduction

‘Cellular plasticity’ is generally defined as the ability of a cell to adapt to different identities along a phenotypic spectrum (*1*). Over the years, plasticity has been extensively studied in neurologic diseases and cancer, to understand the structural and functional changes that can increase adaptability or promote disease progression (*1–3*). Epithelial cells, arranged as cohesive sheets, line body surfaces for specialized roles in absorption, secretion or barrier function. The airway epithelia act as a physical barrier between the external and internal environment and adapts to exposure of more than 10,000 liters of air daily, along with the additional pollutants in the airstream and therefore is an ideal model to understand epithelial responses to chronic injury. A normal pseudostratified airway epithelium undergoes cellular proliferation and differentiation to contribute to functional phenotypes such as permeability, ciliary beat frequency (CBF), and polarity that are required in maintaining tissue homeostasis. Exposure to an irritant or injury can disrupt homeostasis, but plasticity is pivotal in the resilience of the epithelia, permitting them to survive against hostile environment.

In our study, we quantify the phenotypic changes that occur in the barrier function of the airway epithelium. Specifically, we have identified a series of measures of healthy airway epithelium, namely monolayer permeability, mucociliary clearance efficacy, cellular motion or jamming of the epithelial monolayer, cellular polarity, and expression of cell adhesion proteins. The integrity of the airway epithelium is maintained by the ionic gradient and a mechanical barrier (*4, 5*) and we have previously shown that repetitive CS exposure disrupts this integrity (*6*). Persistent exposure to CS results in injury and loss of ciliated cells or cilia (*7, 8*) and a gradual decrease in CBF (*9*), which may lead to remodeling of the epithelia and alteration of mucociliary function (*10–12*). In addition, the establishment of apicobasal epithelial polarity is critical in regulating tissue organization and cellular function (*13, 14*). The most extreme example of epithelial plasticity involves loss of polarity where cells to lose their epithelial identity as occurs with the epithelial to mesenchymal (EMT) transition. EMT requires a loss of cell-cell adhesion, apical-basal polarity and cell motility (*15*). Cell migration is defined as the migratory activity of individual cells. A controlled cell migration mechanism is influenced by cell contraction, expansion and adhesion. A confluent epithelial monolayer does not migrate, however epithelial cells may migrate in response to an injury (*16*). The initial reports of transition between an unjammed to jammed state in the cells in tissues was in published in 2009 (*17*). Additional evidence suggests that epithelial cells from donors with asthma exhibit a delay in the transition to a jammed state (*18*).

Chronic obstructive pulmonary disease (COPD) is characterized by progressive decline in lung function that can occur even after the injurious agent (often tobacco smoke) is removed. It is the third leading cause of death worldwide and fourth leading cause of mortality in United States (*19, 20*). Although dysfunction of several pathways has been implicated in COPD, none of these have resulted in a therapeutic strategy to modify disease. By quantifying the epithelial dysfunction resulting from noxious exposures, such as cigarette smoke (CS), we propose that we can better characterize the stepwise changes that occur from exposure, which we can then compare to phenotypes expressed by cells derived from patients with COPD (COPD human bronchial epithelial; CHBE). Moreover, using pathway inhibitors, we can determine the contribution of several pathways that have been implicated in disease to the epithelial phenotype, and can identify differential behavior in CHBE for better cellular characterization of the patient specific phenotypes.

## Results

### Disrupted epithelial barrier function and adherens junction proteins in CHBE

Although grown under identical conditions, as we have found the pseudostratified age-matched CHBE epithelium formed a leakier monolayer, as measured by a transepithelial electrical resistance (TEER) that is 50% lower and tenfold increase in FITC-dextran flux (Fig. 1A) (*6*). The percentage of ciliated cells (% pixels) was reduced by 60%, and those ciliated cells had a lower CBF, although showed significant donor-specific variability, in CHBE than NHBE (Fig. 1B & C). Moreover, immunohistochemistry of sections of pseudostratified airway epithelium showed delayed differentiation *in vitro* compared to cells derived from normal subjects (Fig. 1D). The CHBE epithelial monolayer was disorganized compared to NHBE, and while there was still evidence of apical-basal polarization based on PKC-ζ and Na,K-ATPase distribution, they exhibited a shorter monolayer height (Fig. 1E).

**Fig. 1.**
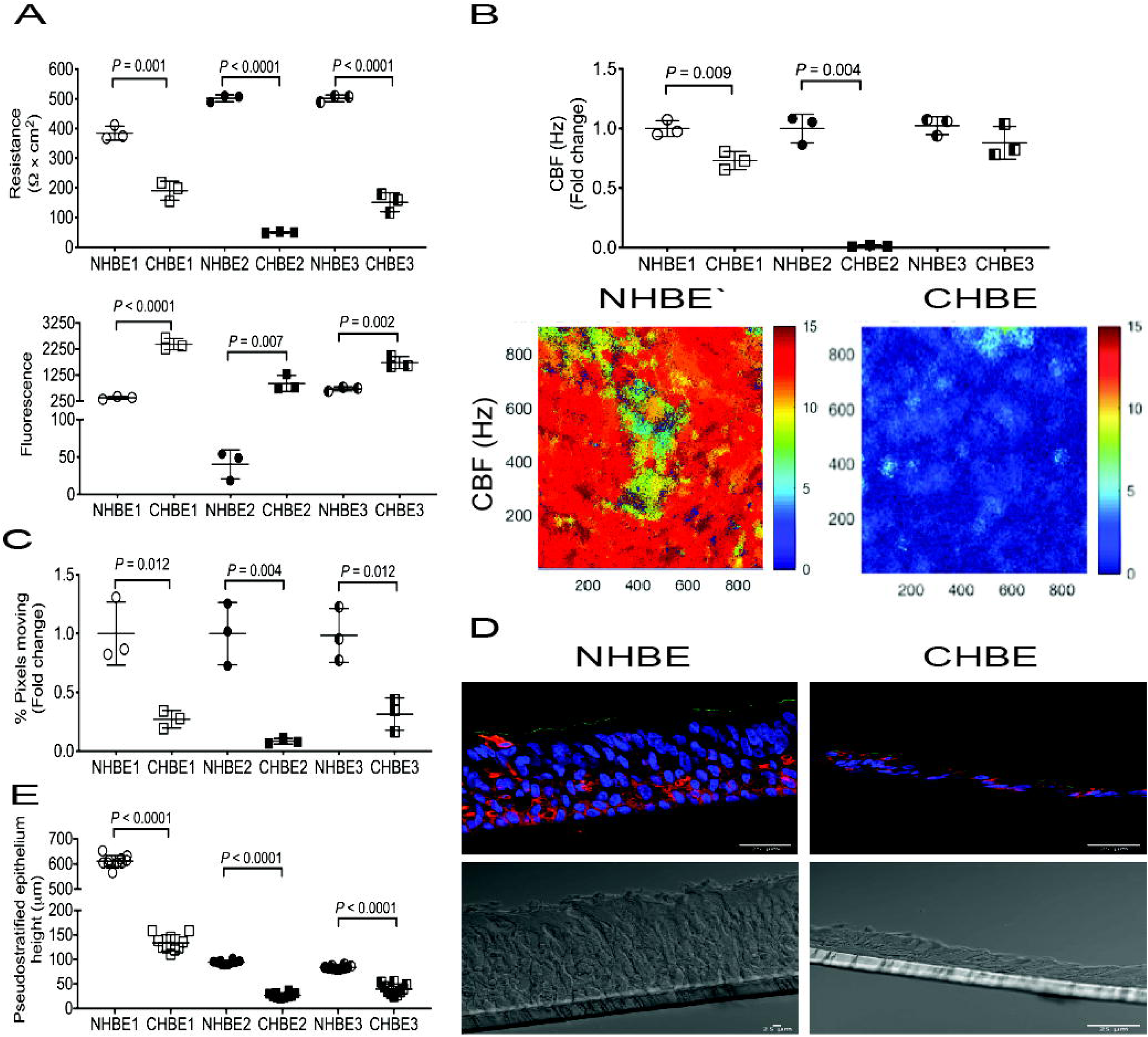
Disruption in barrier function and epithelial polarity in COPD. **(A)** TEER & FITC-Dextran permeability assay, **(B)** Ciliary beat frequency and representative heat maps of CBF, **(C)** % Pixels, **(D)** Immunofluorescence staining of Na, K – ATPase (red) and PKC-ζ (green) and **(E)** Quantified height of pseudostratified epithelium. Images are representative of 3 donors per group (age and gender matched and 3 inserts per donor). Data is expressed as mean ± SD. Shapiro-Wilk normality test followed by Student’s t-test was performed. *P*<0.05 was considered statistically significant. [*Abbreviations*: CBF, ciliary beat frequency; CHBE, COPD derived human bronchial epithelial cells; FITC-Dextran, Fluorescein Isothiocyanate – Dextran; NHBE, Normal human bronchial epithelial cells; TEER, Trans epithelial electrical resistance]

To determine if CHBE demonstrated molecular evidence of cellular plasticity, we measured a series of EMT markers. Compared to NHBE, the mRNA expression of CDH1 was significantly lower in cells derived from CHBE (Fig. S1A). In contrast, the mRNA expression of other EMT markers such as CDH2, VIM, SNAl1, SNAl2, TWIST2, ZEB1, and ZEB2 were higher in CHBE as compared to NHBE (Fig. S1 B-H). To determine if the plasticity indicated a transition to a mesenchymal cell or a different epithelial cell, or both, we looked for the expression of epithelial markers suggestive of squamous metaplasia. In fact, we found that ∼8.5 fold higher levels of p53 in CHBE as compared to NHBE, and although there was a trend towards increase in involcrulin (IVL) was not found to be statistical significant, due to donor variability (Fig. S1 I-J) suggestive that the epithelium was not only demonstrating mesenchymal markers but also squamous cell markers which is in line with the decrease in monolayer height. Also, as we have demonstrated (*6*), the protein expression of E-cadherin was significantly decreased in CHBE compared to NHBE (Fig. S2). The quantitative changes in the phenotype and transcriptional changes suggest evidence of persistent epithelial plasticity in CHBE, although not all of the changes are in line with mesenchymal transition.

#### Effect of EMT-inducer on the barrier function of the primary bronchial epithelia

To evaluate, whether primary bronchial epithelia can undergo EMT transition *in vitro*, the pseudostratified epithelia from healthy donors were treated with 2X EMT-inducer supplement (EMT Supp, R&D Systems), which resulted in a leakier, shorter monolayer (Fig. 2 A, D & E). Although, we observed that cells treated with EMT Supp had lower percentage of ciliated cells, the existing (albeit few) cilia maintained a similar CBF (Figure 2B &C). The decrease in ciliary number was confirmed by scanning electron microscopy (SEM) (Fig. 2F), but the numbers were too low to capture the cross-section of the cilia by transmission electron microscopy (TEM) to confirm the normal 9+2 microtubule arrangement that is normally seen (Fig. 2G).

**Fig. 2.**
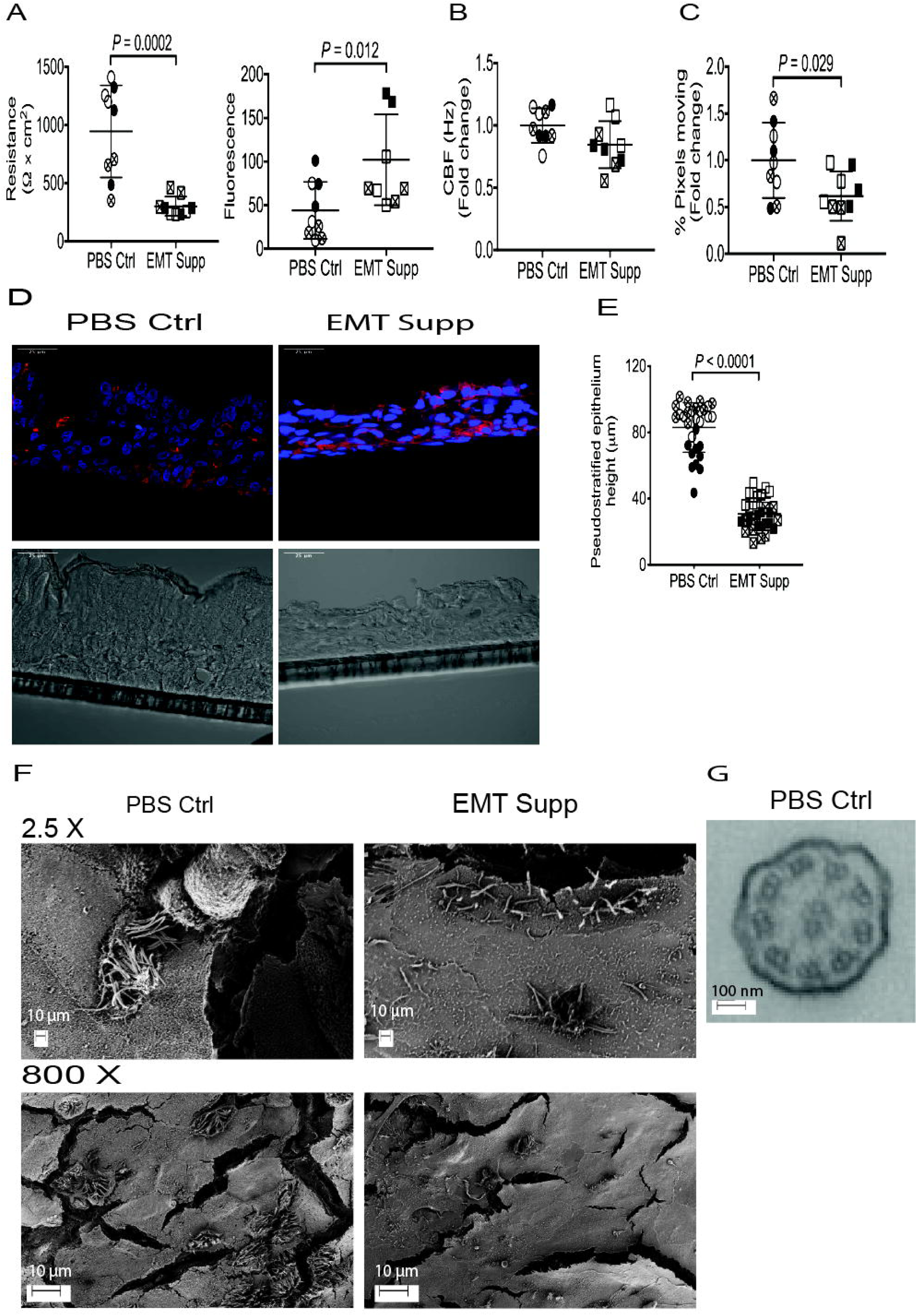
Effect of EMT-inducer on the barrier function of the primary bronchial epithelia. **(A)** TEER & FITC-Dextran permeability assay, **(B)** Ciliary beat frequency, **(C)** % Pixels, **(D)** Immunofluorescence staining of Na, K – ATPase (red), **(E)** Quantified height of pseudostratified epithelium, **(F)** Scanning electron microscopy at 2.5 X and 800 X of the NHBE treated with PBS Ctrl and EMT Supp, and **(G)** Transmission electron microscopy of PBS Ctrl. Images are representative of 3 donors (3 inserts per donor). Data is expressed as mean ± SD. Shapiro-Wilk normality test followed by Student’s t-test was performed. *P*<0.05 was considered statistically significant. [*Abbreviations*: CBF, ciliary beat frequency; EMT Supp, 2X epithelial to mesenchymal transition supplement; FITC-Dextran, Fluorescein Isothiocyanate – Dextran; NHBE, Normal human bronchial epithelia; PBS, phosphate buffer saline, TEER, Trans epithelial electrical resistance]

The NHBE treated with EMT Supp had significantly lower mRNA expression of CDH1 and higher expression of both EMT markers and squamous cell metaplasia markers than the PBS control-treated cells indicating that airway epithelial plasticity involved both transition to both mesenchymal and squamous cells (Fig. 3 A-H). The protein expression of E-cadherin was correspondingly decreased in EMT Supp-treated epithelia compared to PBS control-treated (Fig. S3).

**Fig. 3.**
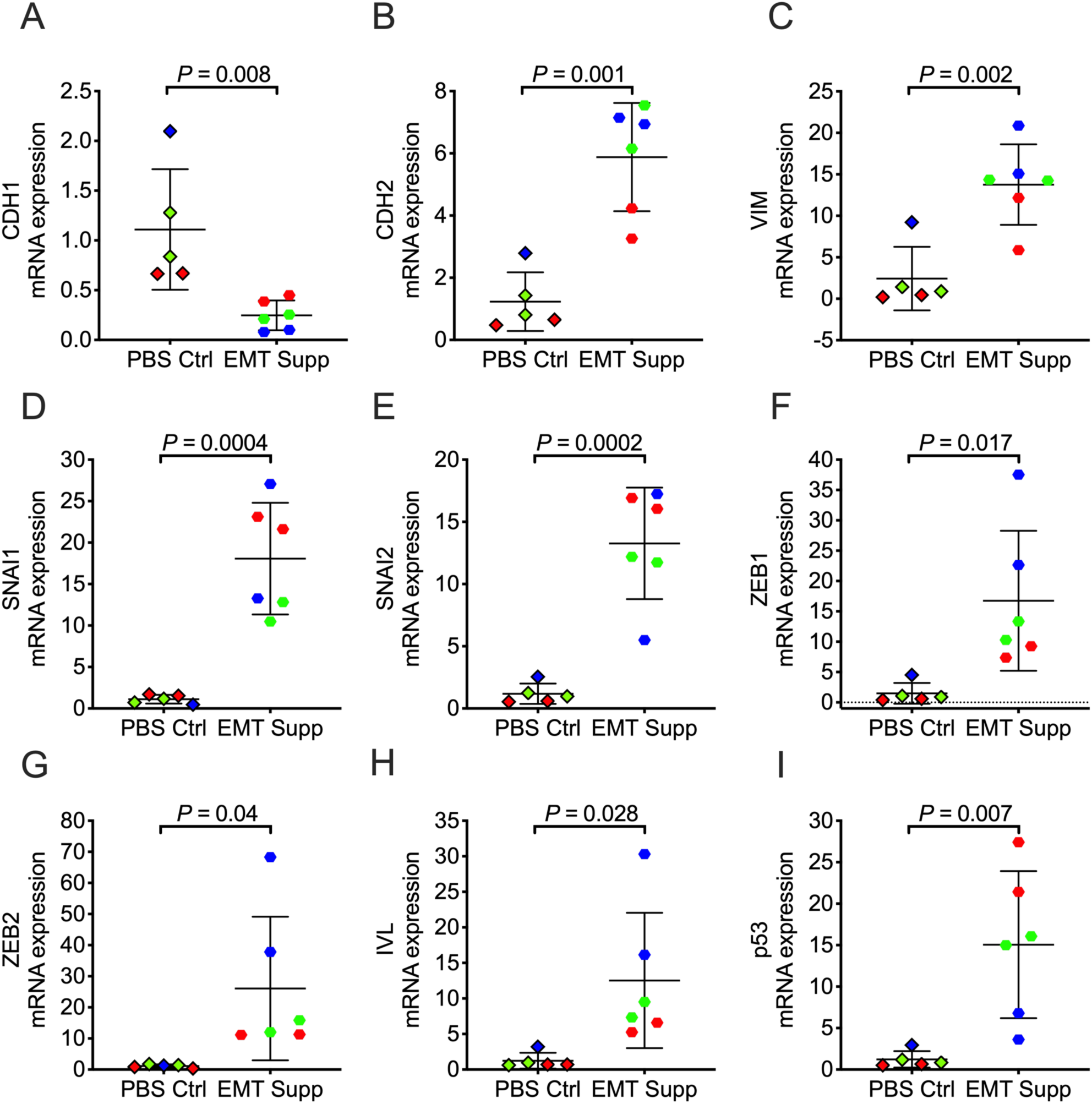
Increased mRNA expression of EMT and squamous markers in EMT-inducing media supplements in healthy epithelia. Basal mRNA expression of epithelial (CDH1) **(A)**, mesenchymal markers (CDH2, VIM, SNAl1, SNAl2, ZEB1 and ZEB2) **(B – G)** and squamous-metaplasia related (p53, IVL) **(H, I)** markers relative to GAPDH in EMT-inducing media supplements compared to PBS in NHBE cells as analyzed by real-time PCR. (n = 3 subjects per group, 2 to 3 inserts per group). Results are shown as mean ±SD. Shapiro-Wilk normality test followed by Student’s t-test was performed. *P*<0.05 was considered statistically significant. [*Abbreviations*: NHBE, normal human bronchial epithelial cells; EMT, Epithelial to mesenchymal transition; PBS, phosphate buffer saline; SD, standard deviation]

#### Epithelial plasticity in cigarette-smoke (CS) exposed NHBE

To determine if CS exposure resulted in similar changes to the epithelial phenotype as seen with the EMT supplement, we quantified the epithelial plasticity. The CS-exposed NHBE formed a leakier monolayer with a 40% decrease in trans-epithelial electrical resistance (TEER) and a tenfold increase in FITC-dextran permeability, reduction in CBF by 18% and had a 40% loss of cilia (as measured by pixels moving) relative to air control (Fig. 4 A-C). Interestingly, repetitive exposure to CS caused a reduction in height of the pseudostratified monolayer compared to air control (Fig. 4D & E). These quantitative phenotypes approached the changes seen in CHBE. The SEM of CS exposed epithelia showed curling of cilia compared to air control, however, there was no difference in 9 + 2 arrangement of microtubules as seen by TEM (Fig. 4F & G). Also, repetitive exposure to CS contributed in significantly lower mRNA expression of CDH1 and higher expression of EMT markers and squamous cell metaplasia markers than the clean air exposed epithelia (Fig. 5 A-I). The expression of EMT markers and the change in monolayer properties are similar to that seen with EMT, although there is no loss in epithelial polarity, showing evidence of persistence of the epithelial phenotype.

**Fig. 4.**
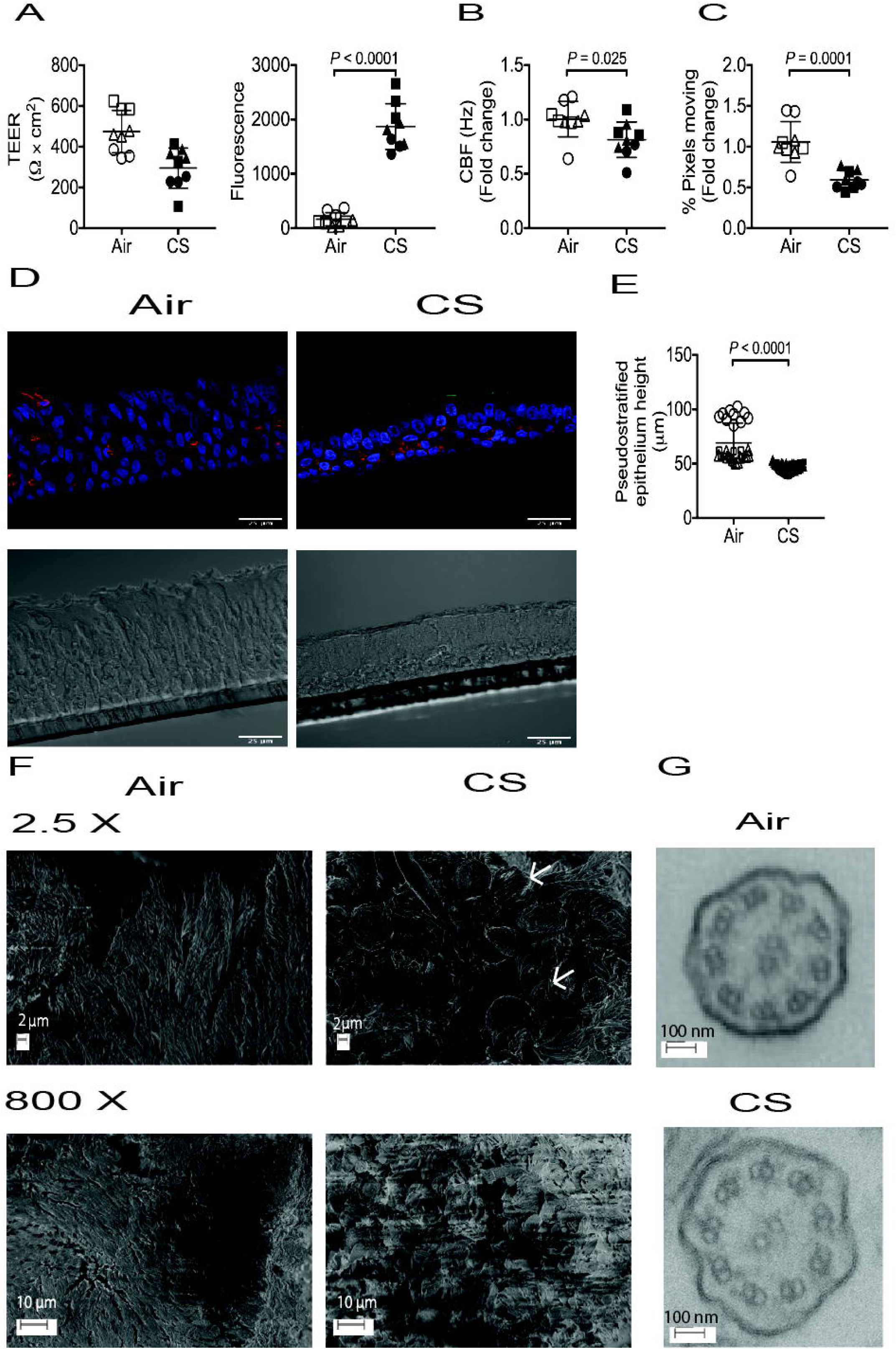
Epithelial plasticity due to repetitive cigarette-smoke exposure. **(A)** TEER & FITC-Dextran permeability assay, **(B)** Ciliary beat frequency, **(C)** % Pixels, **(D)** Immunofluorescence staining of Na, K – ATPase (red) and PKC-ζ (green); **(E)** Height of the NHBE treated with clean air or CS; **(F)** Scanning electron microscopy at 2.5 X and 800 X of the pseudostratified epithelia exposed to CS and clean air control; and **(G)** Transmission electron microscopy of cilia from the NHBE exposed to CS and clean air control. Images are representative of 3 donors (3 inserts per donor). Data is expressed as mean ± SD. Shapiro-Wilk normality test followed by Student’s t-test was performed. *P*<0.05 was considered statistically significant. [*Abbreviations*: CBF, ciliary beat frequency; CS, Cigarette-smoke; FITC-Dextran, Fluorescein Isothiocyanate – Dextran; NHBE, Normal human bronchial epithelia; TEER, Trans epithelial electrical resistance]

**Fig. 5.**
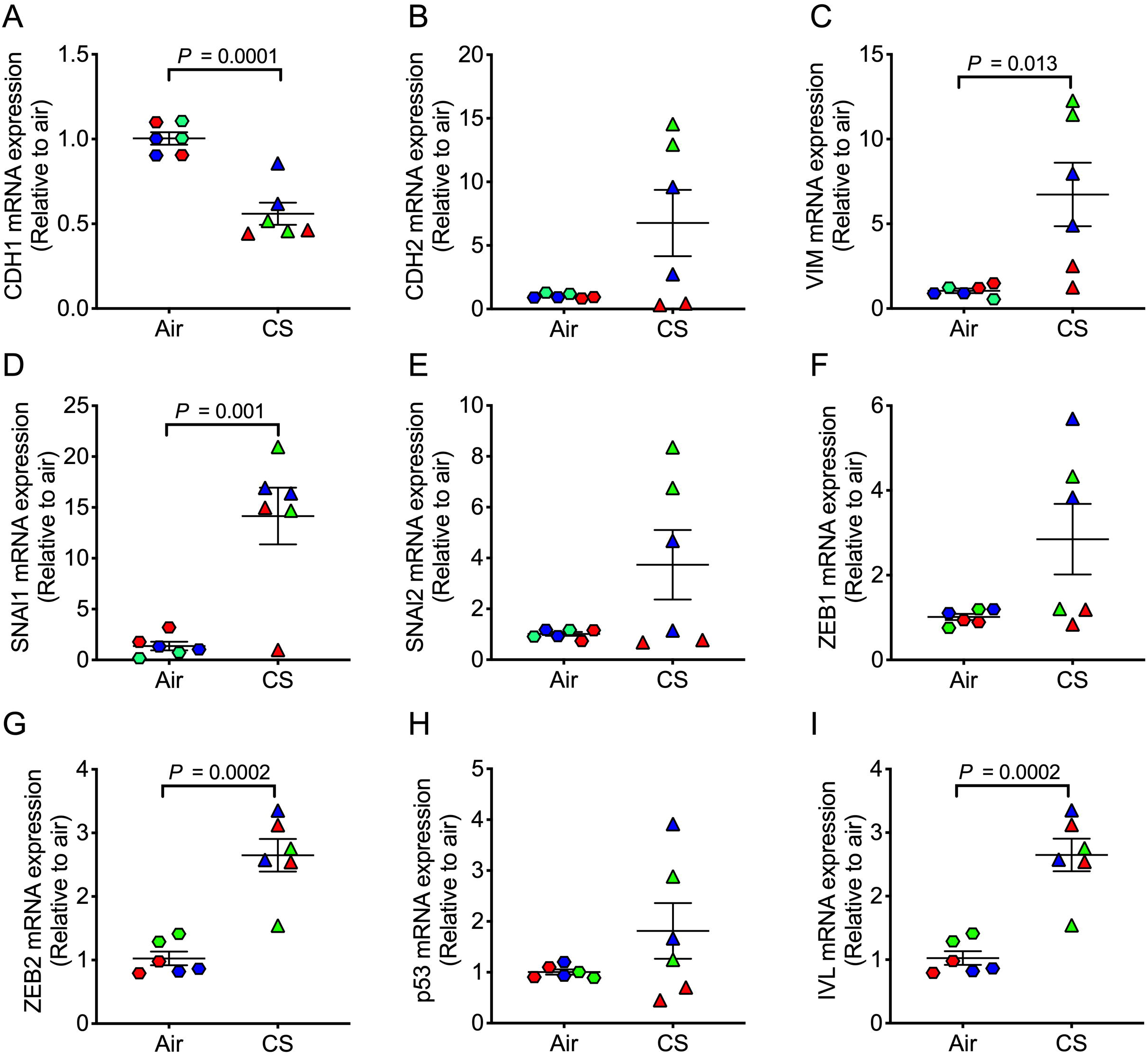
Increased mRNA expression of EMT and squamous markers in cigarette-smoke exposed epithelia. Basal mRNA expression of epithelial (CDH1) **(A)**, mesenchymal markers (CDH2, VIM, SNAl1, SNAl2, ZEB1 and ZEB2) **(B – G)** and squamous-metaplasia related (p53, IVL) **(H, I)** markers relative to GAPDH in cigarette-smoke exposed epithelia compared to clean air exposed epithelia as analyzed by real-time PCR. (n = 3 subjects per group, 2 to 3 inserts per group). Results are shown as mean ±SD. Shapiro-Wilk normality test followed by Student’s t-test was performed. *P*<0.05 was considered statistically significant. [*Abbreviations*: CS, cigarette-smoke; EMT, Epithelial to mesenchymal transition; SD, standard deviation]

#### Jammed to unjammed effect of CS

To further understand the effects of the epithelial plasticity seen on cellular function in the monolayer, we quantified the cellular mobility. Human airway epithelial cells transition from a solid-like jammed state to a more fluid-like unjammed state with the application of compressive stresses on the monolayer (*18*). Moreover, cells derived from asthmatic epithelium have a delayed transition to formation of a jammed state (*18*). We found that the disrupted monolayer and epithelial disorganization seen in CHBE were associated with either a delay in the formation of jammed-like state, or in some patients, a complete lack of jamming, consistent with the variability in the COPD phenotype (Fig. 6A & D). To further elaborate on the dynamics of cooperative cellular mobility, we have calculated the spatial autocorrelation function C(r), from which we estimated the correlation length. The negative values of the correlation function were indicative of swirl patterns in the cell motion, and the correlation length represented a distance over which the cell motion was collective. The correlation length of COPD cells was greater than that seen with NHBE (Fig. 6E), indicating larger intercellular coordination of movement and could be seen visually by prominent swirling motions (Supplementary video). Epithelial plasticity, either due to the EMT supplement (Fig. 6B) or with CS exposure (Fig. 6C) also caused the epithelium to become more fluid-like and unjammed with an increase in correlation length (Fig. 6E) further attesting to cells moving together in a coordinated manner. These data indicated that chemical stimuli can cause the cells transition to a collective movement demonstrating in a fluid-like behavior, even in the absence of changes in cell density, indicating a change in cell shape or stiffness.

**Fig. 6.**
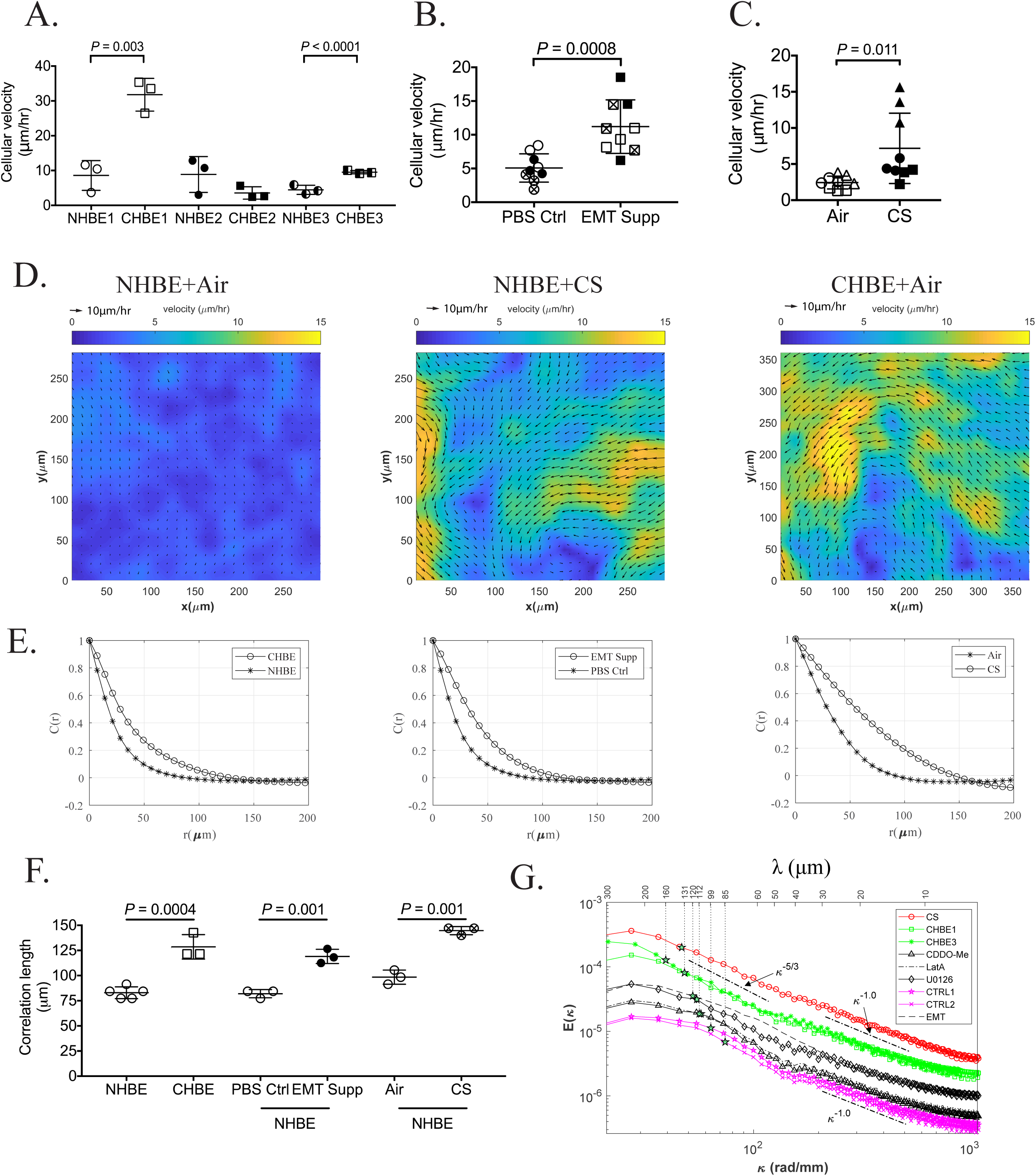
Cell velocity – a proposed marker for epithelial plasticity. Comparison of mean cell velocity between: **(A)** age and gender matched NHBE and CHBE epithelia, **(B)** NHBE treated with PBS Ctrl and EMT Supp, **(C)** Clean air and CS exposed NHBE. Representative heat maps of cell velocity NHBE exposed to air and CS and CHBE exposed to clean air for 10 days **(D).** Distribution of spatial autocorrelation function among NHBE and CHBE epithelia, NHBE treated with PBS Ctrl and EMT Supp, and Clean air and CS exposed NHBE **(E)**. Comparison of correlation length among NHBE and CHBE epithelia, NHBE treated with PBS Ctrl and EMT Supp, and Clean air and CS exposed NHBE **(F)**. Images are representative of 3 donors (3 inserts per donor). Ensembled average one-dimensional spatial energy spectra (E(k), k being wavenumber) of the velocity magnitude **(G)**. The mean length scale or correlation length indicated for each case (stars) can be obtained either from the weighted average of the energy spectrum or from the correlation function. Data is expressed as mean ± SD for cell velocity and correlation length. Shapiro-Wilk normality test followed by Student’s t-test was performed. *P*<0.05 was considered statistically significant. [*Abbreviations*: CHBE, COPD derived human bronchial epithelial cells; CS, cigarette smoke; EMT Supp, 2X epithelial to mesenchymal transition supplement; NHBE, Normal human bronchial epithelial cells; SD, standard deviation]

Figure 6D shows that the flow patterns within the culture consists of multiple swirling eddies of varying sizes and magnitudes. The following analysis attempts to elucidate the size and energy of these eddies for the various cases involved. As a convenient approach, which takes advantage of the wealth of information about typical hydrodynamic motions, we calculated the one-dimensional spatial energy spectra of the velocity component in the same (x) direction (procedures are described in the methods section) for each horizontal line of vectors and then averaged them in time and over the y direction. The energy spectra E(κ), shown in Figure 6G, reflected the distribution of the horizontal contribution to the kinetic energy for varying wavenumbers (κ=2π/λ) or wavelengths (λ), namely the size of eddies. For convenience, results are presented as a function of both wavelength and wavenumber. The relative strength of eddies of different sizes could then be compared to those observed e.g. in two-dimensional (2D) turbulent flows. Note that the above-mentioned correlations were by definition the inverse

Fourier Transform of these spectra, hence corresponding values were also indicated in Figure 6G for each case. These spectra revealed several trends. First, the differences in energy among cases occurred for all eddy sizes, and the energy decreased with decreasing wavelength. Second, the weighted average wavelengths (*i.e.* correlation length) for the CS and two CHBE cases were higher compared to those of the other samples. In contrast, the control cases had the lowest correlation length, and their energy levels were an order of magnitude lower than those of the CHBE and CS cultures. Third, the two CHBE spectra were very similar for most of the spectral range, except for the largest eddies, *i.e.* the differences in velocity magnitude and correlation were associated with the largest scale motions, whereas for smaller scales, the flow patterns were quite similar.

To interpret the differences between jammed to unjammed cultures, we studied the interactions among eddies of different scales in turbulent flow (*21, 22*). These interactions were typically referred to as energy cascading, referring to the transfer of energy from eddies of a certain size to larger or smaller ones owing to merging or breakup, respectively. Unlike 3D flows, in 2D turbulent flows, energy cascading typically occurs in both directions. Upward cascading from small to larger scales owing to merging of eddies occurs in the spectral range corresponding to relatively large eddies. The energy distribution in this spectral range follows the Kolmogorov-Kraichnan scaling, E(κ)=K_ε_ε^2/3^κ^-5/3^, where K_ε_ is a constant and ε is the rate of energy transfer across scales. Such a distribution indicates that the energy flux is scale-independent and stationary, namely the rate at which energy is added to a certain size range due to merging of smaller eddies (ε) is equal to the rate at which the energy is depleted from this range due to merging into larger ones. This energy is eventually dissipated at the largest possible scales owing to interactions e.g. with outer boundaries. In contrast, downward cascading, namely breakup of eddies to smaller ones, occurs at scales that are smaller than those at which the energy is injected. In this range, a stationary down-cascading rate is expected to follow E(κ)∼ κ^-3^ (*22*).

As indicated by lines plotted parallel to the spectra, in the present experiments, spectral ranges with E(κ)∼κ^-5/3^ existed only for the CS and two CHBE cases for λ>40 μm, which corresponded to cell patches with a diameters larger than 5 cells. This trend suggested that the evolution of eddies in the 2D cultures behaved in a similar manner, i.e. the merging into larger ones. The spectral overlap in the κ^-5/3^ domain for the two CHBE cases suggested that they started with very similar energy levels as characteristic cell patches with λ∼40 μm, corresponding to a diameter of about 5 cells, and had the same energy flux (merging rate) up to 160 um (∼25 cells diameter). However, for some reason, in one of the cases, CHBE3, the cascading process persisted to larger eddies compared to that observed for CHBE1. Such differences could be associated with variations in e.g. friction at the interface with the microscope or along the outer perimeter (*23*). Yet, for most of the spectral range, the distribution and energy of the eddy motions for the two CHBE cultures were very similar. The CS flow also followed the reverse cascading process, but the initial energy level, at λ∼40 μm as well, and the subsequent kinetic energy flux, were about two times higher. As discussed later, this difference could be associated with an increase in the stresses initiating the motions at the scales at which the energy was injected or a decrease in the strength of adherens junctions between cells, which would reduce their ability to withstand these stresses. In contrast, lack of a E(κ)∼κ^-5/3^ range for the control cases and cultures treated with drugs indicated that their initial kinetic energy levels, also injected at λ∼40 μm (suggested from the kinks in the spectra), were too weak to initiate a stationary reverse cascading process (merging) of eddies. As discussed later, this lower initial kinetic energy could have been caused by decreases in the initial stresses and/or cell stiffness as well as by an increase in the strength of adherens junctions, i.e. an improved ability to withstand these stresses.

#### Effect of inhibitors and activator on barrier function

Except for a narrow range of the CDDO-Me case, the present cultures did not exhibit an E(κ)∼κ^-3^ range at scales smaller than that of the injected energy, which is expected for stationary breakup to small scale eddies(*22*). Instead, in almost all cases, the small-scale motions exhibited E(κ)∼κ^-1^. For 2D flows, direct transition from E(κ)∼κ^-5/3^ to κ^-1^ is typically observed in spectra of passive scalars (temperature or mass diffusion) where the mass diffusivity is much smaller than the viscosity (*24, 25*). In the E(κ)∼κ^-1^ range, the viscosity dampened the velocity fluctuations, but the diffusivity is too small to smoothen the scalar fluctuations. Instead, the spatial variability in scalar concentration is progressively smeared by weak stresses induced by residual motions. Making an analogy to the present cell cultures, the direct transition from E(κ)∼κ^-5/3^ to κ^-1^ suggested that once the initial stresses fragmented the cultures to patches with characteristic size of 5 cells diameter, part of the energy cascaded to larger scales. For the remaining energy, the residual stresses within each sub-patch were too weak to cause further fragmentation. Consequently, the slow energy transfer across scales causing the κ^-1^ domain could have been a result of e.g. residual stresses within each patch or stresses induced by relative motion between patches.

To evaluate if CS induced disruption in barrier function can be blocked, we treated the CS exposed epithelium with specific inhibitors of actin polymerization (LatA), of MAPK/ERK kinase (U0126), Nrf-2 pathway activation (CDDO-Me), TGFβ1 neutralizer (1D11) and antagonist of Wnt signaling pathway (Dkk1). We have demonstrated that after repetitive exposure to CS, there is increased cortical tension in the epithelium, and therefore included low dose latrunculin to cause some actin depolymerization to decrease cortical tension, while not killing the cells (*6*). In fact, we observed that Lat A, U0126 and CDDO-Me exhibited a protective effect against repetitive CS exposure by inhibiting the CS induced leak (Fig. 7A) and decreases in CBF (Fig. 7B). However, these inhibitors had no effect in restoring the cilia (Fig. 7C). Also, the same three drugs (Lat A, U0126 and CDDO-Me) were found to be protective against jamming to unjamming transition (Fig. 7D), with decreases in the correlation length scale induced by CS (Fig. 7E) suggestive that the same mechanisms that mediate the disruption in barrier integrity are also involved in the transition to an unjammed state involve similar mechanisms. Interestingly, CDDO-Me was found to protective against CS induced suppression of CDH1 expression, indicating its protective role (Fig. 8A).

**Fig. 7.**
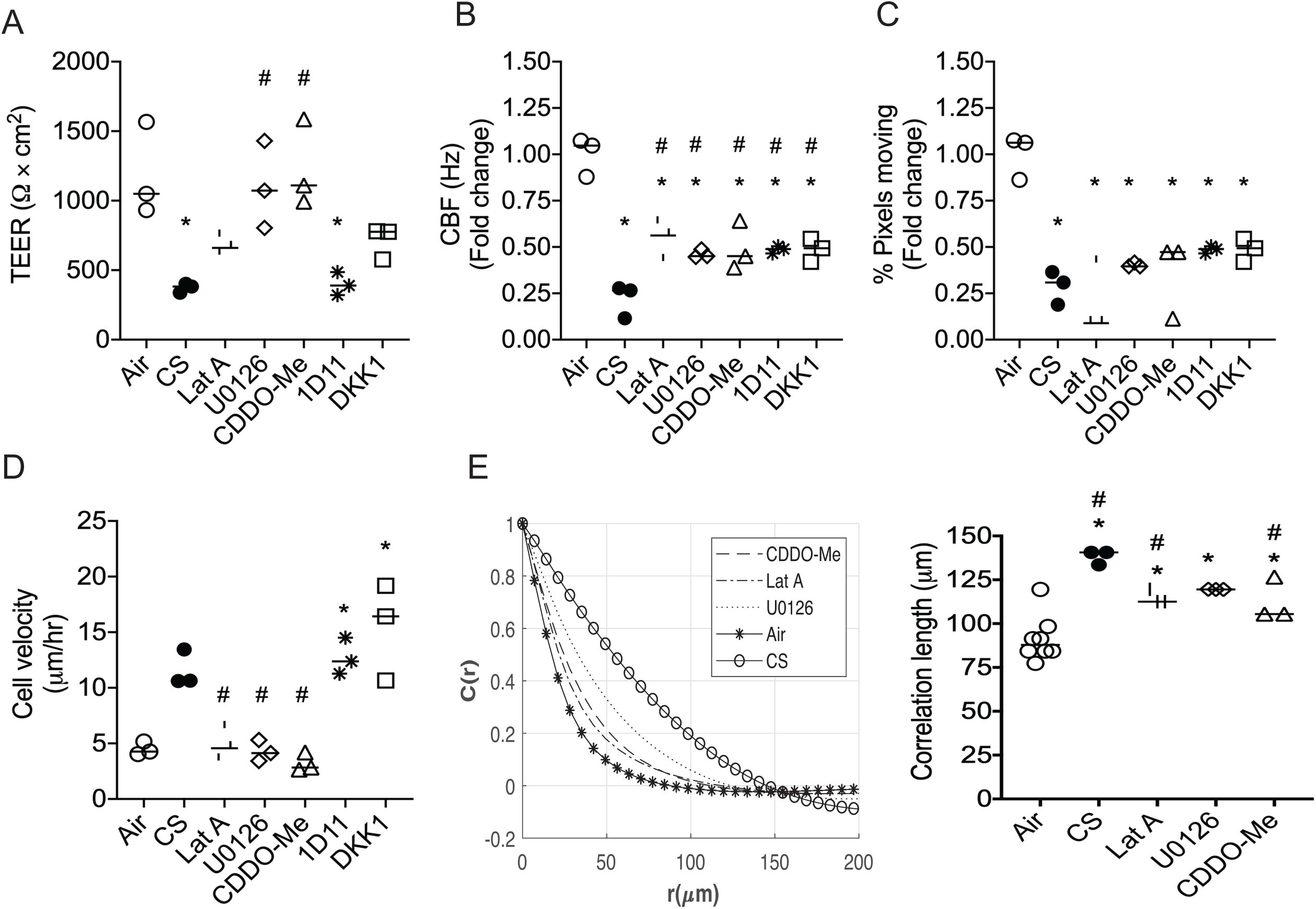
Modulating the epithelial plasticity by inhibiting specific pathways. Effect of specific pathway inhibitors on TEER **(A)**, Ciliary beat frequency **(B)**, % Pixels **(C)**, and cell velocity **(D)**. Effect of inhibitors on the distribution of spatial correlation function, Correlation length **(E)** and mRNA expression of CDH1 **(F)**. Images are representative of 3 inserts from one donor. Epithelia were treated with inhibitor of actin polymerization (Lat A), MAPK kinase inhibitor (U0126), Nrf2 activator (CDDO-Me), TGFβ1 neutralizer (1D11); and antagonist of Wnt signaling pathway (DKK1). Data is expressed as median. Shapiro-Wilk normality test followed by Student’s t-test was performed. \**P*<0.05 was considered statistically significant. [*Abbreviations*: CBF, ciliary beat frequency; CDDO-Me, CDDO-Methyl ester; CS, cigarette-smoke; Lat A, Latrunculin A; TEER, Trans epithelial electrical resistance]

**Fig. 8.**
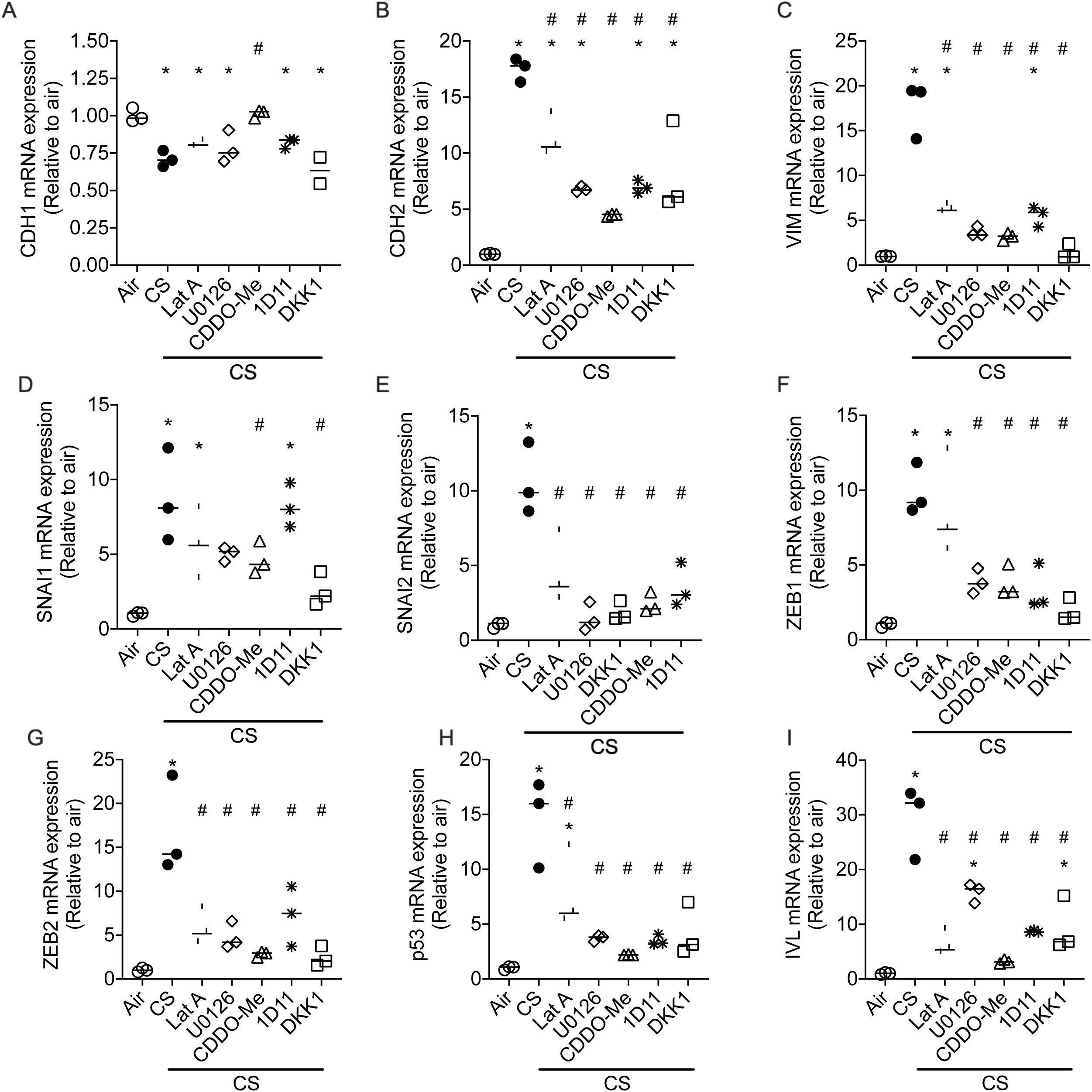
Effect of pathway inhibitors and activators on cigarette-smoke exposed epithelia. Basal mRNA expression of epithelial (CDH1) **(A)**, mesenchymal markers (CDH2, VIM, SNAl1, SNAl2, ZEB1 and ZEB2) **(B – G)** and squamous-metaplasia related (p53, IVL) **(H, I)** markers relative to GAPDH in cigarette-smoke exposed epithelia compared to clean air exposed epithelia as analyzed by real-time PCR. Data is expressed as median from 3 inserts from one donor. Shapiro-Wilk normality test followed by Student’s t-test was performed. *P*<0.05 was considered statistically significant. [*Abbreviations*: CS, cigarette-smoke; EMT, Epithelial to mesenchymal transition; SD, standard deviation]

## Discussion

Cellular plasticity, or the ability of the cell to adapt both structurally and functionally, is critical to cellular resilience in the face of chronic injury. This study shows has shown that repetitive exposure to CS induces a quantitative phenotypic change in the airway epithelium, which could be extended to other injurious stimuli. We observe that even in the absence of injurious substance, the cells often cannot revert back to the non-diseased phenotype, as in the context of COPD airway epithelium.

In this paper, we have quantified the phenotypic changes that occur in the epithelium in CHBE cells and CS exposed NHBE cells, and have determined that these changes represent plasticity along the spectrum of epithelial-to-mesenchymal transition (EMT), by comparing it to cells treated with an EMT-inducing supplement. We found that although the COPD epithelium maintained cell polarity, and therefore did not completely transition to mesenchymal cells, there was evidence of plasticity based on poor monolayer integrity, monolayer shortening and disorganization, decreased CBF and numbers of ciliated cells, as well as increased kinetic energy, causing a fluid-like behavior of the monolayer (*6*)(*26*)(*27*). Along with the down-regulation of E-cadherin mRNA and protein, we also found that transcriptional markers seen in more mesenchymal cells as well as in squamous epithelium were upregulated in repetitive CS-exposed non-diseased epithelia, indicating that CS induces a similar phenotypic shift to that of the COPD epithelia.

Transitioning to the unjammed state has been proposed as an alternative strategy to EMT for cellular mobility (*28*), and we observe transition into the unjammed state does occur with epithelial plasticity albeit without the requirement of loss of apical-basal polarity. This transition occurs without any change in cellular density, indicating that the increased motion is not simply due to an uncaging of the cell but due to altered cellular mechanics in response to the chemical injury in the case of CS, or persistent alterations in mechanical properties in CHBE cells. Based on our current and previous data (*6*) we have found that exposure to cigarette smoke has two relevant mechanical effects – it increases the cell stiffness and decreases cell-cell contact strength. There is conflicting literature on the correlation between cell stiffness and migration (*29*), with some studies showing that the cell motility improves with decreased stiffness (*30*), but others indicating that the stiffer cells are more prone to become motile (*31*). Yet, since the increased stiffness reduces the cell’s ability to deform, once the culture is exposed to some stresses, the loading on the adherens junction between the more rigid cells should increase. Owing to the accompanying reduction in cell-cell contact strength, the adherens junctions are more likely to break up, causing fragmentation of the culture into migrating patches.

Consequently, while we do not know whether the cigarette smoke affects the external forces driving the culture dynamics, even with similar levels of forcing, the much higher initial kinetic energy of the CS culture is consistent with the increased stiffness and reduced contact strength. To alleviate these effects, one could either decrease the cell stiffness, achieved in the present study using LatA, which decreases actin polymerization, or enhance the cell-cell contact strength by treating the culture with CDDO-Me, which as we show is the only compound to increase E-cadherin levels in our study. As the present results show, both cause a drastic reduction in the initial kinetic energy and subsequent culture migration. This could also be aided by the fact that latrunculin also it reduces new actin filament growth, which is necessary for motility. The kinetic energy spectra of cell-patch migration indicate that for the CHBE and CS-exposed cultures, the initial kinetic energy injected as the culture is fragmented into patches, with a characteristic diameter of about 5 cells, is high enough to cause behavior analogous to that of a two-dimensional turbulent fluid flow. Included are a spectral range with −5/3 slope, suggesting merging (reverse cascading) of cell patches into larger scale eddies, and a weak/slow breakup into smaller eddies. The (2x) higher initial kinetic energy and merging rate for the CS case compared to that of CHBE could be associated with higher cell stiffness, weaker adherens junctions among cells, or possibly, differences in the initial forcing. The two CHBE cases show similar initial and cascading rate of energy, with the differences between them occurring only at the scales where the large-scale energy is dissipated. In contrast, owing to the lower stiffness and/or stronger adherens junctions for the control and drug-treated CS-exposed cases, the initial kinetic energy levels are too weak to generate multiscale motions analogous to turbulent flows. Yet, even after drug treatment, the CS-exposed cultures are still more energetic than the controls.

Comparing other pathway inhibitors implicated in COPD and with EMT (*32–36*), we studied manipulating mitogen-activated protein kinase (MAPK)/extracellular-signal-regulated kinase (ERK), Nrf-2, TGFβ, and Wnt signalling pathway, with improvement in the identified barrier properties seen only with inhibition of actin polymerization (LatA), activation of the Nrf-2 pathway (CDDO-Me) and MAPK/ERK kinase (U0126), although none of these completely restored ciliary mechanics. Whether manipulation of all of these represent mechanisms to alter cortical tension and/or adhesion between cells or contribute to other compensatory pathways need further study. Our data suggests that a therapeutic strategy is to target both reduction of cellular stresses and increase of junctional stresses. We propose that chronic low-grade injury causes the epithelium to undergo plasticity to provide tissue resilience but result in phenotypic changes that alter diminishes homeostatic functions. Quantification of the phenotype from a given patient can provide a strategy to personalize therapeutic strategies given the diversity in the clinical presentation as well as provides an assessment of reversal that occurs either with individual therapies or with combinations of drugs.

## Materials and Methods

NHBE were either treated with EMT inducer supplement or were exposed to whole CS in an exposure system and VC 1 manual smoking machine (Vitrocell^®^ Systems GmbH, Germany) as described previously (*6*). We confirmed the changes in the epithelial plasticity phenotypes such as permeability by trans-epithelial electrical resistance (TEER) and fluorescein isothiocyanate-dextran (FITC-Dextran) flux assay, ciliary beat frequency (CBF), cell migration (detailed protocol described below) and polarity by immunofluorescence (SI appendix). To verify any changes in the structure of cilia, samples were collected for scanning electron microscope (SEM) and transmission electron microscope (TEM) (SI appendix). The protein expression of E-cadherin and EMT markers were quantified by western blotting and RT-qPCR respectively (SI appendix).

### Cell migration

For quantifying cell migration, phase contrast images of cell culture were acquired with one frame taken every 5 minutes for 2 hours (as described in CBF). The successive images were cross-correlated in a similar manner to standard Particle Image velocimetry using the commercial software DaVis (LaVison Inc., MI, USA) (*37*). Multi-pass cross-correlation analysis with decreasing interrogation window size were used for optimizing the data quality. The size of the final correlated windows was 32 × 32 pixels, corresponding to 5.4 µm × 5.4 µm. With a 50% overlap between the neighboring windows, the resulting spatial resolution of the vector field was 2.7 µm × 2.7 µm. Vector data post-processing was performed by universal outlier detection to remove any spurious vectors (*38*). For each data set, the instantaneous velocities were computed by averaging the instantaneous quantities over the entire field of view. All the instantaneous quantities were time-averaged over a period of 2 hours. To quantify the intercellular coordination, the spatial autocorrelation function (*39*) was determined at each time point *t* as

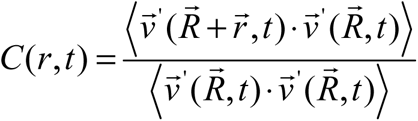

Where 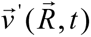 denotes the velocity fluctuations at the position 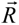, obtained by subtracting the spatial mean from the velocity field, and angular brackets indicate an average over all positions 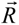. The correlation function was then averaged over all directions and times. The distance where the correlation function first decays to zero was taken as the correlation length (*40*).

The spatial one-dimensional energy spectra of the velocity component in x-direction, E(κ), was calculated from the instantaneous velocity field along a series of horizontal lines (*41, 42*). The fast Fourier transform of the velocity field along the horizontal lines was computed after detrending and subtracting the spatially averaged velocity from each line, with-out any windowing functions. The instantaneous spectrum obtained for the multiple lines and times were then ensemble averaged to obtain the mean spatial energy spectra. Note that the correlation is the inverse Fourier Transform of the spectrum, i.e. it represents the average length scale of the cell culture motions.

## Acknowledgement

We would like to thank Johns Hopkins University School of Medicine Microscope Facility for providing access to 3i Marianis Spinning Disk Confocal, and Zeiss LSM700 single-point laser scanning confocal microscope. We would like to thank Reynold A. Panettieri (Director, Rutgers Institute for Translational Medicine and Science, New Brunswick, New Jersey) for providing for providing age, and gender matched NHBE and CHBE cells. We wish to thank Pablo A. Iglesias (Electrical and Computer Engineering, Johns Hopkins Whiting School of Engineering, Baltimore, Maryland) for designing the MatLab script to analyse the CBF of the pseudostratified epithelium.

Research reported in this publication was supported by the National Heart, Lung, and Blood Institute (NHLBI & R01-HL124099) and Office of the Director of the National Institutes of Health under award number S10OD016374.

This research was made possible in part by a grant from The Gulf of Mexico Research Initiative. Data are publicly available through the Gulf of Mexico Research Initiative Information & Data Cooperative (GRIIDC) at https://data.gulfresearchinitiative.org (https://doi.org/10.7266/n7-xpky-ss66).

## Supplementary Materials and Methods

### Primary human airway epithelial cells

Primary human bronchial epithelial cells from healthy (NHBE) and COPD donors were either sourced from MatTek Corporation (MA, United States), Lonza Group AG (Basel, Switzerland) or Epithelix SàRL (Geneva, Switzerland). Demographic characteristics of the donors are shown in Table 1.

### Cell culture

The cryopreserved cells were amplified on Collagen I Rat tail (Corning®, NY, USA) coated flasks with PneumaCult^TM^-Ex Plus medium (StemCell Technologies Inc., Vancouver, Canada). Cells were passaged at 80-90% confluency. At sub-confluency cells were plated onto Rat tail Collagen I coated 0.4 μm pore polyethylene terephthalate clear membrane transwell inserts (Corning®, NY, USA) with PneumaCult^TM^-Ex Plus medium at air-liquid interface (ALI) with 150,000 cells/well, 400,000 cells/well and 750,000 cells/well of 24-well, 12-well, and 6-well transwells respectively. At 100% confluency, the transwells were put at air-liquid interface (ALI) with basolateral PneumaCult^TM^-ALI medium (StemCell Technologies Inc., Vancouver, Canada). Cells were differentiated for 4 to 6 weeks ALI to obtain a fully differentiated pseudostratified epithelium.

### Cigarette-smoke (CS) exposure

For whole CS experiments, the confluent monolayer of pseudostratified epithelium was exposed to CS in an exposure system and VC 1 manual smoking machine (Vitrocell^®^ Systems GmbH, Germany) as described previously (*6*). Once CS exposure consisted of 2 cigarettes (3R4F) and each burned for ∼8 minutes using the ISO puff regimen (One 35 mL puff every 60 seconds with an 8 second exhaust). The control inserts were exposed to humidified air in the exposure system.

### Trans-epithelial electrical resistance (TEER) measurement

On the apical compartment of the inserts, sterile 1X phosphate buffered saline (PBS, ThermoFisher Scientific, NY, USA) was added. TEER was then assessed three times with an epithelial voltohmmeter (EVOM, World Precision Instruments, FL, USA) with the STX2 electrodes of 4 mm width and 1 mm thick and a mean resistance was calculated. Values were then corrected for fluid resistance (insert with no cells) and surface area.

### Permeability

Permeability was assessed by fluorescein isothiocyanate-dextran (FITC-dextran) flux assay as described previously (*6*). Briefly 1mg/mL of 4 kDA FITC-dextran (Sigma-Aldrich, MO, USA) diluted in 1X PBS was added to the apical surface of monolayer with basolateral 1X PBS. After 1 hour, transwells were removed and basolateral 1X PBS were transferred to a clear bottom 96-well plate and fluorescence was measured in triplicate with an excitation of 485 nm and emission of 528 nm in a Synergy^TM^ HT multi-detection microplate reader (Biotek, VT, USA).

### CBF

Differentiated pseudostratified epithelium were placed into No. 1.5 coverslip glass top and bottom plates (MatTek Corporation, MA, USA). The plates were incubated at 37°C with 5% CO_2_ in the 3i Marianis Spinning Disk Confocal-phase contrast microscope system (Leica Microsystems, Wetzlar Germany). High-speed time-lapse images were taken at a magnification of 32x using a scientific CMOS camera (Hamamatsu C11440-42U30, Hamamatsu Photonics K.K., Hamamatsu City, Japan). For measuring CBF, a sequence of 250, 1800×1800 pixels images were acquired at 100 frames per second. In all the cases, the field of view was 304 x 304*µ*m^2^. A Matlab (R2018b, The MathWorks, Inc., MA, USA) program (*43*) was used to determine average CBF per video. Refer to earlier studies (*9*) for more details on CBF analysis.

### Immunofluorescence

Differentiated pseudostratified epithelium were fixed with 4% paraformaldehyde in PBS (Affymetrix Inc, Ohio, US) to the apical chamber and incubated for 15 minutes. All incubations were carried out at room temperature. The cells were washed with 1X PBS and were transferred through a series of sucrose (Sigma-Aldrich, MO, USA) infiltration steps. Following sucrose infiltration, 10 μm vertical sections of the membrane were performed on Leica Cryostat (CM3050 S, IL, USA) and immunostaining for mouse monoclonal sodium/potassium-transporting ATPase subunit alpha-1 (Na+/K+-ATPase α, 1:100, Santa Cruz Biotechnology, Inc) and mouse monoclonal protein kinase C (PKC ζ, 1:50, Santa Cruz Biotechnology) was conducted as described in SI (Materials and Methods). The slides were imaged on Single-point, laser scanning confocal microscope (LSM 700, Carl Zeiss Microscopy LLC, NY, USA) at 40X oil.

### Sucrose infiltration and immunostaining

The sucrose infiltration steps involved addition of each solution in apical and basolateral side of inserts for 10 minutes, 10% sucrose, 2:1 10%:30% sucrose, 1:1 10%:30% sucrose, 1:2 10%:30% sucrose, and 30% sucrose. Following sucrose infiltration, the membrane containing the cells was embedded in a biopsy size cryomold using Optimal Cutting Temperature (O.C.T.) Compound (Tissue-Tek, CA, USA). 10 μm sections were cut on the and attached to Superfrost Plus microscope slides (Fisher Scientific, PA, USA) and dried at room temperature for overnight.

Slides were stained using the Shandon Sequenza Immunostaining center (Fisher Scientific, PA, USA). The sections were permeabilized with 0.1% Triton^TM^ X-100 (Fisher Scientific, NJ, USA) in 1X PBS, blocked at room temperature for 1 to 2 hours with 5% BSA (Bovine Serum Albumin, (Sigma-Aldrich, MO, USA) and 10% goat serum (ThermoFisher Scientific, CA, USA) in 1X PBS, and incubating the primary antibody 1:100 to 1:300 dilution overnight at 4°C. Primary antibody was washed thrice with 1X PBS and secondary antibody 1:200 in 1X PBS were added to each slide and incubated for 3 hrs at room temperature in the dark. The secondary antibody was then washed, and the slides are mounted with ProLong® Gold antifade with DAPI (ThermoFisher Scientific, CA, USA). Slides were cured overnight, sealed, and stored at 4°C for long term storage.

### Transmission Electron Microscopy (TEM)

*Sample preparation:* Samples were fixed in 2.5% glutaraldehyde, 3mM MgCl_2_, in 0.1 M sodium cacodylate buffer (pH 7.2) overnight at 4°C. After buffer rinse, samples were post fixed in 1% osmium tetroxide in 0.1 M sodium cacodylate buffer on ice in the dark for 1 hour. The samples were washed with 0.1 M sodium cacodylate buffer to remove the excess fixative buffer. Samples were incubated at 4°C overnight in filtered sterile 0.1 M sodium cacodylate buffer, washed with 0.1 M maleate buffer, en bloc stained with 2% uranyl acetate (for 1 hr in the dark) in 0.1 M maleate, dehydrated in a graded series of ethanol, propylene oxide and embedded in Eponate 12 (Ted Pella) resin. Samples were polymerized at 60°C overnight. Thin sections, 60 to 90 nm, were cut with a diamond knife on the Reichert-Jung Ultracut E ultramicrotome and picked up with naked 200 mesh copper grids. Grids were stained with 2% uranyl acetate (aq.) followed by lead citrate.

The samples were observed with a Philips CM120 Cryo-Electron Microscope at 80 kV and the images were captured with an AMT XR80 high-resolution (16-bit) 8 Megapixel camera (Advanced Microscopy Techniques, MA, USA).

### Scanning Electron Microscopy (SEM)

*Sample Preparation:* Samples were fixed in 2.5% glutaraldehyde, 3mM MgCl_2_, in 0.1 M sodium cacodylate buffer (pH 7.2) overnight at 4°C. After buffer rinse, samples were post fixed in 1% osmium tetroxide in buffer (1 hr) on ice in the dark followed by two distilled water rinses before dehydration in ethanol. Samples were dried for SEM with hexamethyldisilazane, mounted on carbon coated stubs, and coated with 20 nm gold/palladium alloys (AuPd).

Samples were prepared as describe in SI and imaged on a Leo/Zeiss Field-emission SEM at 1 kV.

### Quantitative reverse transcription polymerase chain reaction (RT-qPCR)

Total RNA was isolated from cultured primary bronchial epithelial cells and purified using the RNeasy® Plus Mini Kit (Qiagen, Germany), supplemented with the Proteinase K (Qiagen, Germany) and RNase-Free Dnase Set (Qiagen, Germany). cDNA of 1000 ng µL^-1^ was obtained using the High Capacity cDNA Reverse Transcription Kit (Applied Biosystems, ThermoFisher Scientific Baltics, Lithuania), and the absence of DNA contamination was verified by excluding the reverse transcriptase from subsequent PCR reactions.

cDNA was subjected to PCR using the SYBR^TM^ Green PCR Master Mix (Applied Biosystems, Thermo Fisher Scientific, UK) to amplify CDH1 (Epithelial-Cadherin), CDH2 (Neural-Cadherin), VIM (Vimentin), SNAl1 (Snail Family Transcriptional Repressor 1), SNAl2 (Snail Family Transcriptional Repressor 2), TWIST1 (Twist-related protein 1), ZEB1 (Zinc Finger E-Box Binding Homobox 1), ZEB2 (Zinc Finger E-Box Binding Homobox 1), p53 (tumour protein), and GAPDH (Glyceraldehyde 3-phosphate dehydrogenase) using primers (integrative DNA Technologies, IA) (Table S2).

Each PCR reaction was carried out as follows: Initial denaturation at 94°C for 15 minutes, 45 cycles of 94°C for 35 seconds, 60°C for 1 minute and 72°C for 1 minute 15 seconds, followed by a final extension at 72°C for 2 minutes. Each cycle was repeated 45 times. Based on comparative Ct method, gene expression levels were calculated and GAPDH was used as housekeeping gene.

### EMT induction in NHBE

To induce EMT in NHBE cells (5-6-week ALI) were additionally supplemented with the basolateral 2X EMT induction supplement (StemXVivo EMT Inducing Media Supplement, R&D Systems) for 10 days.

### Western blotting assay

Whole cells extracts were prepared with 1X RIPA buffer (Cell Signalling Technology®), quantified for total protein using bicinchoninic acid colorimetric assay (Pierce™ BCA Protein Assay Kit, IL, USA), separated on sodium dodecyl sulfate – polyacrylamide gel electrophoresis (SDS-PAGE) using the 4-15% Mini-PROTEAN® TGX™ Precast Protein Gels and Mini-PROTEAN^®^ Tetra Vertical Electrophoresis Cell (Bio-Rad), transferred to PVDF (Immobilon^®^-FL PVDF membrane, Millipore Sigma, County Cork, Ireland) using Mini Trans-Blot^®^ Cell (Bio-Rad), and then incubated with primary monoclonal antibody [E-cadherin (24E10) Rabbit mAb (1:1000, Cell Signaling Technology®) or β-actin (13E5) Rabbit mAb (1:1000, Cell Signaling Technology®)] and probed with secondary antibody [1:20,000, IRDye^®^ 800CW Donkey anti-Rabbit IgG (H+L) or IRDye^®^ 680CW Goat anti-Rabbit IgG (H+L)]. The blots were imaged by ODYSSEY CLx imaging system and quantified using Image Studio Lite (LI-COR Biosciences).

### Drug treatments

The NHBE cells at ALI were exposed to CS for 10 days and simultaneously basolateral treated with an antagonist of Wnt signaling pathway – 50ng/mL of Recombinant basolateral treated with an antagonist of Wnt signaling pathway – 50ng/mL of Recombinant Human Dkk-1 Protein (R&D Systems) or TGFβ1 neutralizer – 1.25 µg/mL of TGF beta-1,2,3 Monoclonal Antibody (1D11) (ThermoFisher Scientific) or MAPK kinase inhibitor – 15μM of U0126 (Cell Signaling Technology®) or Nrf2 activator – 50nM of CDDO-Me (Toronto Research Chemicals) or an inhibitor of actin polymerization – 0.625μM of Latrunculin A (Sigma-Aldrich).

### Statistical analysis

Prism version 7.0 (GraphPad, CA, USA) was used for analysis. The results are expressed as a mean ± SD or median. A *P*<0.05 was considered statistically significant. Comparisons between two groups were performed using an unpaired Student’s *t*-test. One-way ANOVA with Tukey’s multiple comparison test were performed to evaluate the effect of pathway inhibitors or activators on barrier function.

**Table S1:** Demographic characteristics of donors

**A. Donor characteristics for age and gender matched donors**
| **Characteristics** | **NHBE1** | **CHBE1** | **NHBE2** | **CHBE2** | **NHBE3** | **CHBE3** |
| --- | --- | --- | --- | --- | --- | --- |
| **Age (in years)** | 69 | 67 | 77 | 79 | 71 | 64 |
| **Gender** | Male | Male | Male | Male | Female | Female |
| **Smoking history** | -- | Ex-smoker<br>(25 pack-years) | -- | Ex-smoker<br>(20 pack-years) | Yes* | Yes* |

**B. Donor characteristics for healthy epithelia treated with EMT inducing supplement or exposed to air or cigarette-smoke**
| **Characteristics** | **NHBE** |
| --- | --- |
| **N** | 4 |
| **Age (in years)** | 55.25 ± 6.70 |
| **Gender (Male : Female)** | 1 : 3 |
| **Smoking history**<br>**(Non-smoker : Smoker)** | 2 : 2* |

\*Pack-years data not available.
Age is shown as mean ±SD.
[NHBE, normal human bronchial epithelial cells; CHBE, COPD-derived human bronchial epithelial cells; SD, standard deviation]

**Table S2:**
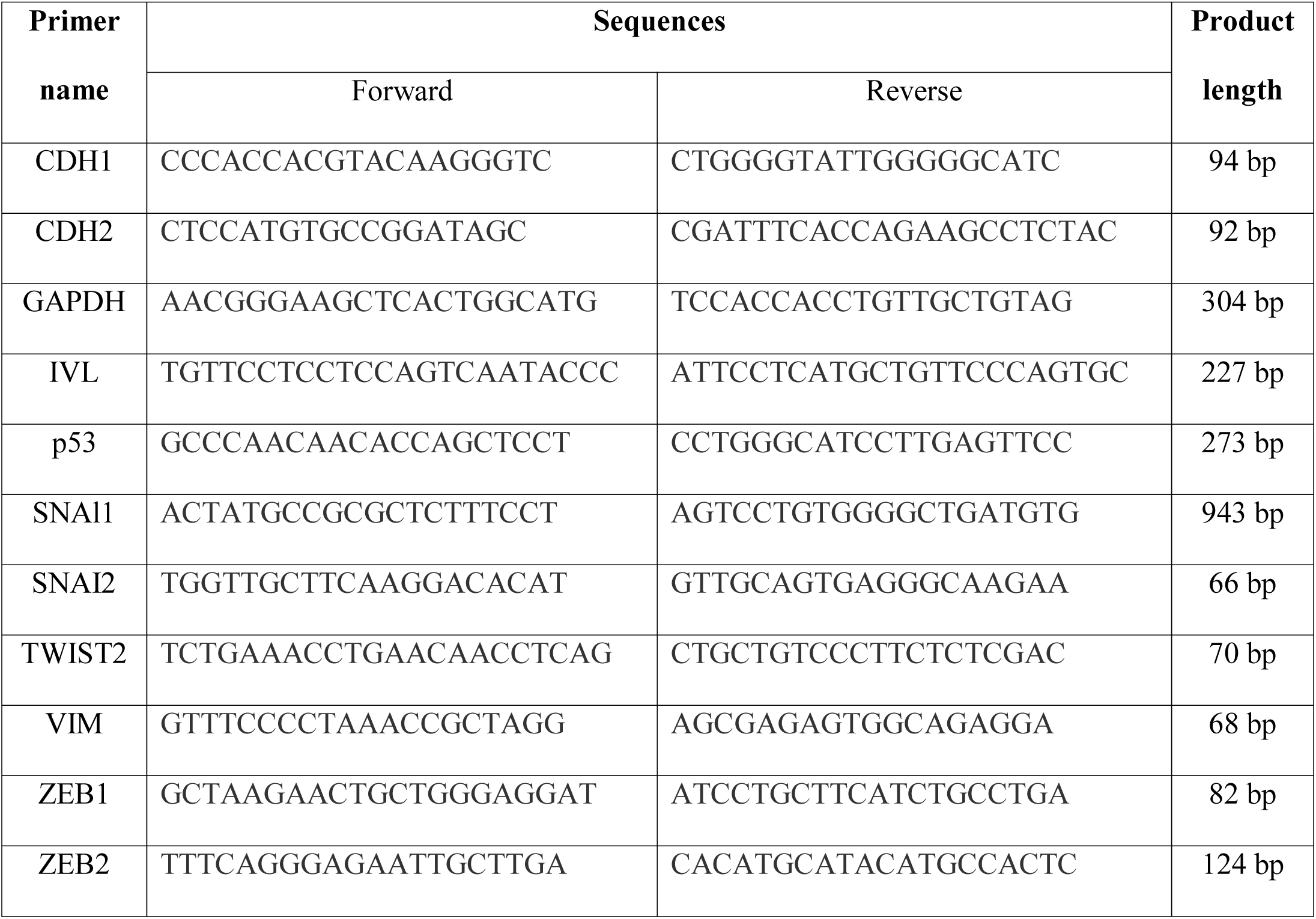
Primers for quantitative PCR analysis of gene transcript expression

## Figures

**Fig. S1. Reduced mRNA expression of epithelial markers and increased mRNA expression of EMT- and squamous-metaplasia related markers in COPD subjects.** Basal mRNA expression of epithelial (CDH1) (A), mesenchymal markers (CDH2, VIM, SNAl1, SNAl2, ZEB1 and ZEB2) (B – H) and squamous-metaplasia related (p53, IVL) (I, J) markers relative to GAPDH in cigarette-smoke exposed epithelia compared to clean air exposed epithelia as analyzed by real-time PCR. (n = 3 subjects per group, 1 to 3 inserts per group). Results are shown as mean ±SD. Shapiro-Wilk normality test followed by Student’s t-test was performed. *P*<0.05 was considered statistically significant. [NHBE, normal human bronchial epithelial cells; CHBE, COPD-derived human bronchial epithelial cells; EMT, Epithelial to mesenchymal transition; SD, standard deviation]

**Fig. S2. Reduced expression of E-cadherin protein in COPD subjects.** Representative blots of E-cadherin expression from NHBE and CHBE cells as detected by western blotting. β-Actin levels are shown as a loading control (n = 3 subjects per group, 1 to 2 inserts per group). Results are shown as mean ±SD. Shapiro-Wilk normality test followed by Student’s t-test was performed. *P*<0.05 was considered statistically significant. [NHBE, normal human bronchial epithelial cells; CHBE, COPD-derived human bronchial epithelial cells; SD, standard deviation]

**Fig. S3. Reduced expression of E-cadherin protein in EMT Supp treated NHBE.** β-Actin levels are shown as a loading control (n = 2 subjects per group, 2 inserts per donor). Results are shown as mean ±SD. Shapiro-Wilk normality test followed by Student’s t-test was performed. *P*<0.05 was considered statistically significant. [NHBE, normal human bronchial epithelial cells; SD, standard deviation]

## Movies

**S1:** Representative movie showing the cell migration for NHBE exposed to air for 10 days.

**S2:** Representative movie showing the cell migration for NHBE exposed to cigarette-smoke for 10 days.

**S3:** Representative movie showing the cell migration for CHBE exposed to air for 10 days.

## References

1. S. Yuan, R. J. Norgard, B. Z. Stanger, Cellular Plasticity in Cancer, Cancer Discov. (2019), doi:10.1158/2159-8290.CD-19-0015.

2. B. Kolb, R. Gibb, Brain plasticity and behaviour in the developing brain, J. Can. Acad. Child Adolesc. Psychiatry 20, 265–276 (2011).

3. A. Surcel, E. S. Schiffhauer, D. G. Thomas, Q. Zhu, K. T. DiNapoli, M. Herbig, O. Otto, H. West-Foyle, A. Jacobi, M. Krater, K. Plak, J. Guck, E. M. Jaffee, P. A. Iglesias, R. A. Anders, D. N. Robinson, Targeting mechanoresponsive proteins in pancreatic cancer: 4-hydroxyacetophenone blocks dissemination and invasion by activating MYH14, Cancer Res. 79, 4665–4678 (2019).

4. M. J. Welsh, Electrolyte transport by airway epithelia, Physiol. Rev. 67, 1143–1184 (2017).

5. S. Ganesan, A. T. Comstock, U. S. Sajjan, Barrier function of airway tract epithelium, Tissue Barriers 1, e24997 (2013).

6. K. Nishida, K. A. Brune, N. Putcha, P. Mandke, W. K. O’Neal, D. Shade, V. Srivastava, M. Wang, H. Lam, S. S. An, M. B. Drummond, N. N. Hansel, D. N. Robinson, V. K. Sidhaye, Cigarette smoke disrupts monolayer integrity by altering epithelial cell-cell adhesion and cortical tension, Am. J. Physiol. Cell. Mol. Physiol. (2017), doi:10.1152/ajplung.00074.2017.

7. A. Livraghi, S. H. Randell, Cystic Fibrosis and Other Respiratory Diseases of Impaired Mucus Clearance, Toxicol. Pathol. 35, 116–129 (2007).

8. K. S. Park, J. M. Wells, A. M. Zorn, S. E. Wert, V. E. Laubach, L. G. Fernandez, J. A. Whitsett, Transdifferentiation of ciliated cells during repair of the respiratory epithelium, Am. J. Respir. Cell Mol. Biol. 34, 151–157 (2006).

9. L. D. Chandrala, N. Afshar-Mohajer, K. Nishida, Y. Ronzhes, V. K. Sidhaye, K. Koehler, J. Katz, A Device for measuring the in-situ response of Human Bronchial Epithelial Cells to airborne environmental agents, Sci. Rep. 9, 7263 (2019).

10. B. Thomas, A. Rutman, C. O’Callaghan, Disrupted ciliated epithelium shows slower ciliary beat frequency and increased dyskinesia, Eur. Respir. J. 34, 401–404 (2008).

11. A. Yaghi, A. Zaman, G. Cox, M. B. Dolovich, Ciliary beating is depressed in nasal cilia from chronic obstructive pulmonary disease subjects, Respir. Med. 106, 1139–1147 (2012).

12. B. Thomas, A. Rutman, R. A. Hirst, P. Haldar, A. J. Wardlaw, J. Bankart, C. E. Brightling, C. O’Callaghan, Ciliary dysfunction and ultrastructural abnormalities are features of severe asthma, J. Allergy Clin. Immunol. 126, 722–729.e2 (2010).

13. L. M. McCaffrey, I. G. Macara, Widely conserved signaling pathways in the establishment of cell polarity., Cold Spring Harb. Perspect. Biol. 1, 1–17 (2009).

14. P. D. Wilson, Epithelial cell polarity and disease, Am. J. Physiol. Physiol. 272, F434–F442 (1997).

15. J. P. Thiery, Epithelial-mesenchymal transitions in development and pathologies, Curr. Opin. Cell Biol. 15, 740–746 (2003).

16. M. Poujade, A. Hertzog, J. Jouanneau, P. Chavrier, B. Ladoux, A. Buguin, P. Silberzan, Collective migration of an epithelial monolayer, Proc. Natl. Acad. Sci. 104, 15988–15993 (2007).

17. X. Trepat, M. R. Wasserman, T. E. Angelini, E. Millet, D. A. Weitz, J. P. Butler, J. J. Fredberg, Physical forces during collective cell migration, Nat. Phys. 5, 426 (2009).

18. J. A. Park, J. H. Kim, D. Bi, J. A. Mitchel, N. T. Qazvini, K. Tantisira, C. Y. Park, M. McGill, S. H. Kim, B. Gweon, J. Notbohm, R. Steward, S. Burger, S. H. Randell, A. T. Kho, D. T. Tambe, C. Hardin, S. A. Shore, E. Israel, D. A. Weitz, D. J. Tschumperlin, E. P. Henske, S. T. Weiss, M. L. Manning, J. P. Butler, J. M. Drazen, J. J. Fredberg, Unjamming and cell shape in the asthmatic airway epithelium, Nat. Mater. (2015), doi:10.1038/nmat4357.

19. R. Lozano, M. Naghavi, K. Foreman, S. Lim, K. Shibuya, V. Aboyans, J. Abraham, T. Adair, R. Aggarwal, S. Y. Ahn, M. A. AlMazroa, M. Alvarado, H. R. Anderson, L. M. Anderson, K. G. Andrews, C. Atkinson, L. M. Baddour, S. Barker-Collo, D. H. Bartels, M. L. Bell, E. J. Benjamin, D. Bennett, K. Bhalla, B. Bikbov, A. Bin Abdulhak, G. Birbeck, F. Blyth, I. Bolliger, S. Boufous, C. Bucello, M. Burch, P. Burney, J. Carapetis, H. Chen, D. Chou, S. S. Chugh, L. E. Coffeng, S. D. Colan, S. Colquhoun, K. E. Colson, J. Condon, M. D. Connor, L. T. Cooper, M. Corriere, M. Cortinovis, K. C. de Vaccaro, W. Couser, B. C. Cowie, M. H. Criqui, M. Cross, K. C. Dabhadkar, N. Dahodwala, D. De Leo, L. Degenhardt, A. Delossantos, J. Denenberg, D. C. Des Jarlais, S. D. Dharmaratne, E. R. Dorsey, T. Driscoll, H. Duber, B. Ebel, P. J. Erwin, P. Espindola, M. Ezzati, V. Feigin, A. D. Flaxman, M. H. Forouzanfar, F. G. R. Fowkes, R. Franklin, M. Fransen, M. K. Freeman, S. E. Gabriel, E. Gakidou, F. Gaspari, R. F. Gillum, D. Gonzalez-Medina, Y. A. Halasa, D. Haring, J. E. Harrison, R. Havmoeller, R. J. Hay, B. Hoen, P. J. Hotez, D. Hoy, K. H. Jacobsen, S. L. James, R. Jasrasaria, S. Jayaraman, N. Johns, G. Karthikeyan, N. Kassebaum, A. Keren, J.-P. Khoo, L. M. Knowlton, O. Kobusingye, A. Koranteng, R. Krishnamurthi, M. Lipnick, S. E. Lipshultz, S. L. Ohno, J. Mabweijano, M. F. MacIntyre, L. Mallinger, L. March, G. B. Marks, R. Marks, A. Matsumori, R. Matzopoulos, B. M. Mayosi, J. H. McAnulty, M. M. McDermott, J. McGrath, Z. A. Memish, G. A. Mensah, T. R. Merriman, C. Michaud, M. Miller, T. R. Miller, C. Mock, A. O. Mocumbi, A. A. Mokdad, A. Moran, K. Mulholland, M. N. Nair, L. Naldi, K. M. V. Narayan, K. Nasseri, P. Norman, M. O’Donnell, S. B. Omer, K. Ortblad, R. Osborne, D. Ozgediz, B. Pahari, J. D. Pandian, A. P. Rivero, R. P. Padilla, F. Perez-Ruiz, N. Perico, D. Phillips, K. Pierce, C. A. Pope, E. Porrini, F. Pourmalek, M. Raju, D. Ranganathan, J. T. Rehm, D. B. Rein, G. Remuzzi, F. P. Rivara, T. Roberts, F. R. De León, L. C. Rosenfeld, L. Rushton, R. L. Sacco, J. A. Salomon, U. Sampson, E. Sanman, D. C. Schwebel, M. Segui-Gomez, D. S. Shepard, D. Singh, J. Singleton, K. Sliwa, E. Smith, A. Steer, J. A. Taylor, B. Thomas, I. M. Tleyjeh, J. A. Towbin, T. Truelsen, E. A. Undurraga, N. Venketasubramanian, L. Vijayakumar, T. Vos, G. R. Wagner, M. Wang, W. Wang, K. Watt, M. A. Weinstock, R. Weintraub, J. D. Wilkinson, A. D. Woolf, S. Wulf, P.-H. Yeh, P. Yip, A. Zabetian, Z.-J. Zheng, A. D. Lopez, C. J. Murray, Global and regional mortality from 235 causes of death for 20 age groups in 1990 and 2010: a systematic analysis for the Global Burden of Disease Study 2010, Lancet 380, 2095–2128 (2012).

20. D. M. Mannino, A. S. Buist, Global burden of COPD : risk factors, prevalence, and future trends, Lancet 370, 765–773 (2007).

21. H. Tennekes, J. Lumley, A First Course in Turbulence, Book (1972), doi:10.1017/S002211207321251X.

22. R. H. Kraichnan, Inertial-range transfer in two-and three-dimensional turbulence, J. Fluid Mech. (1971), doi:10.1017/S0022112071001216.

23. R. H. Kraichnan, Inertial ranges in two-dimensional turbulence, Phys. Fluids (1967), doi:10.1063/1.1762301.

24. K. R. Sreenivasan, Turbulent mixing: A perspective, Proc. Natl. Acad. Sci. (2019), doi:10.1073/pnas.1800463115.

25. T. Gotoh, J. Nagaki, Y. Kaneda, Passive scalar spectrum in the viscous-convective range in two-dimensional steady turbulence, Phys. Fluids (2000), doi:10.1063/1.870291.

26. A. Yaghi, M. Dolovich, Airway Epithelial Cell Cilia and Obstructive Lung Disease, Cells 5, 40 (2016).

27. A. C. Schamberger, N. Mise, J. Jia, E. Genoyer, A. Ö. Yildirim, S. Meiners, O. Eickelberg, Cigarette smoke-induced disruption of bronchial epithelial tight junctions is prevented by transforming growth factor-β, Am. J. Respir. Cell Mol. Biol. 50, 1040–1052 (2014).

28. J.-A. Park, L. Atia, J. A. Mitchel, J. J. Fredberg, J. P. Butler, Collective migration and cell jamming in asthma, cancer and development, J. Cell Sci. 129, 3375–3383 (2016).

29. Q. Luo, D. Kuang, B. Zhang, G. Song, Cell stiffness determined by atomic force microscopy and its correlation with cell motility, Biochim. Biophys. Acta - Gen. Subj. 1860, 1953–1960 (2016).

30. C. Mihai, S. Bao, J. P. Lai, S. N. Ghadiali, D. L. Knoell, PTEN inhibition improves wound healing in lung epithelia through changes in cellular mechanics that enhance migration, Am. J. Physiol. - Lung Cell. Mol. Physiol. (2012), doi:10.1152/ajplung.00037.2011.

31. K. Radotić C. Roduit, J. Simonović, P. Hornitschek, C. Fankhauser, D. Mutavdžić Steinbach, G. Dietler, S. Kasas, Atomic force microscopy stiffness tomography on living arabidopsis thaliana cells reveals the mechanical properties of surface and deep cell-wall layers during growth, Biophys. J. (2012), doi:10.1016/j.bpj.2012.06.046.

32. J. Shankar, I. R. Nabi, Actin cytoskeleton regulation of epithelial mesenchymal transition in metastatic cancer cells, PLoS One 10, 1–12 (2015).

33. L. Xie, B. K. Law, A. M. Chytil, K. A. Brown, M. E. Aakre, H. L. Moses, Activation of the Erk pathway is required for TGF-β 1-induced EMT in vitro, Neoplasia 6, 603–610 (2004).

34. F. Bocci, S. C. Tripathi, S. A. Vilchez Mercedes, J. T. George, J. P. Casabar, P. K. Wong, S. M. Hanash, H. Levine, J. N. Onuchic, M. K. Jolly, NRF2 activates a partial epithelial-mesenchymal transition and is maximally present in a hybrid epithelial/mesenchymal phenotype, Integr. Biol. 11, 251–263 (2019).

35. J. Xu, S. Lamouille, R. Derynck, TGF-β-induced epithelial to mesenchymal transition, Cell Res. 19, 156–172 (2009).

36. K. Gasior, M. Hauck, A. Wilson, S. Bhattacharya, A Theoretical Model of the Wnt Signaling Pathway in the Epithelial Mesenchymal Transition, Theor. Biol. Med. Model. 14, 1–20 (2017).

37. M. Raffael, C. Willert, S. T. Wereley, J. Kompenhans, Particle Image Velocimetry (the Third Edition) (2007).

38. J. Westerweel, F. Scarano, Universal outlier detection for PIV data, Exp. Fluids (2005), doi:10.1007/s00348-005-0016-6.

39. S. Garcia, E. Hannezo, J. Elgeti, J.-F. Joanny, P. Silberzan, N. S. Gov, Physics of active jamming during collective cellular motion in a monolayer, Proc. Natl. Acad. Sci. (2015), doi:10.1073/pnas.1510973112.

40. W. Xi, S. Sonam, T. Beng Saw, B. Ladoux, C. Teck Lim, Emergent patterns of collective cell migration under tubular confinement, Nat. Commun. (2017), doi:10.1038/s41467-017-01390-x.

41. C. Li, J. Miller, J. Wang, S. S. Koley, J. Katz, Size Distribution and Dispersion of Droplets Generated by Impingement of Breaking Waves on Oil Slicks, J. Geophys. Res. Ocean. (2017), doi:10.1002/2017JC013193.

42. E. E. Hackett, L. Luznik, J. Katz, T. R. Osborn, Effect of finite spatial resolution on the turbulent energy spectrum measured in the coastal ocean bottom boundary layer, J. Atmos. Ocean. Technol. (2009), doi:10.1175/2009JTECHO647.1.

43. J. H. Sisson, J. A. Stoner, B. A. Ammons, T. A. Wyatt, All-digital image capture and whole-field analysis of ciliary beat frequency, J. Microsc. 211, 103–111 (2003).

