## Supplementary figures and images for "Quantifying Epithelial Plasticity as a Platform to Reverse Epithelial Injury"

### Figure S1

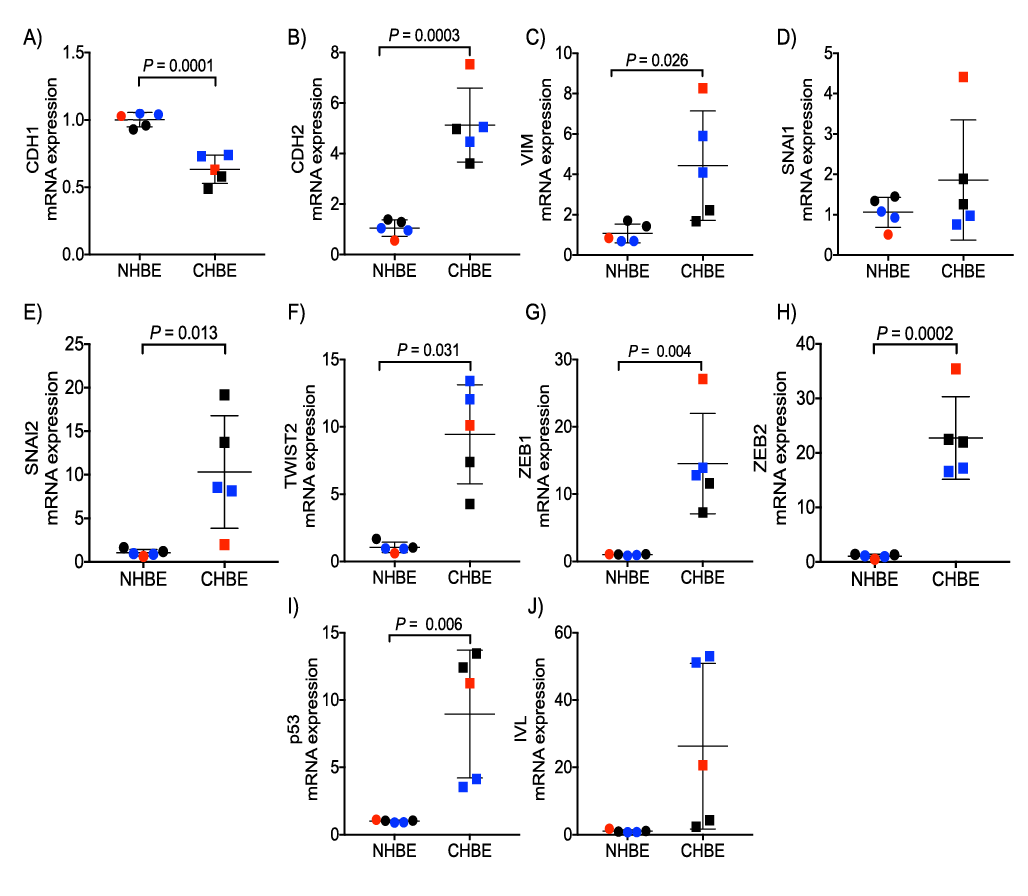

### Figure S2

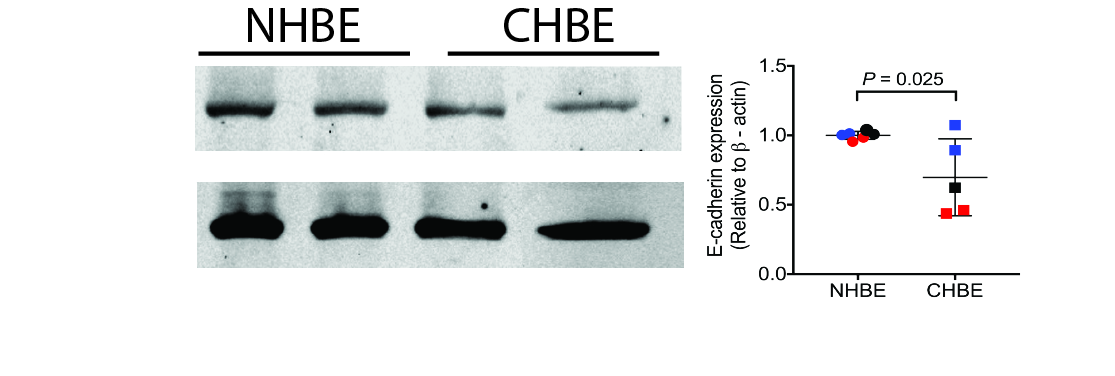

### Figure S3

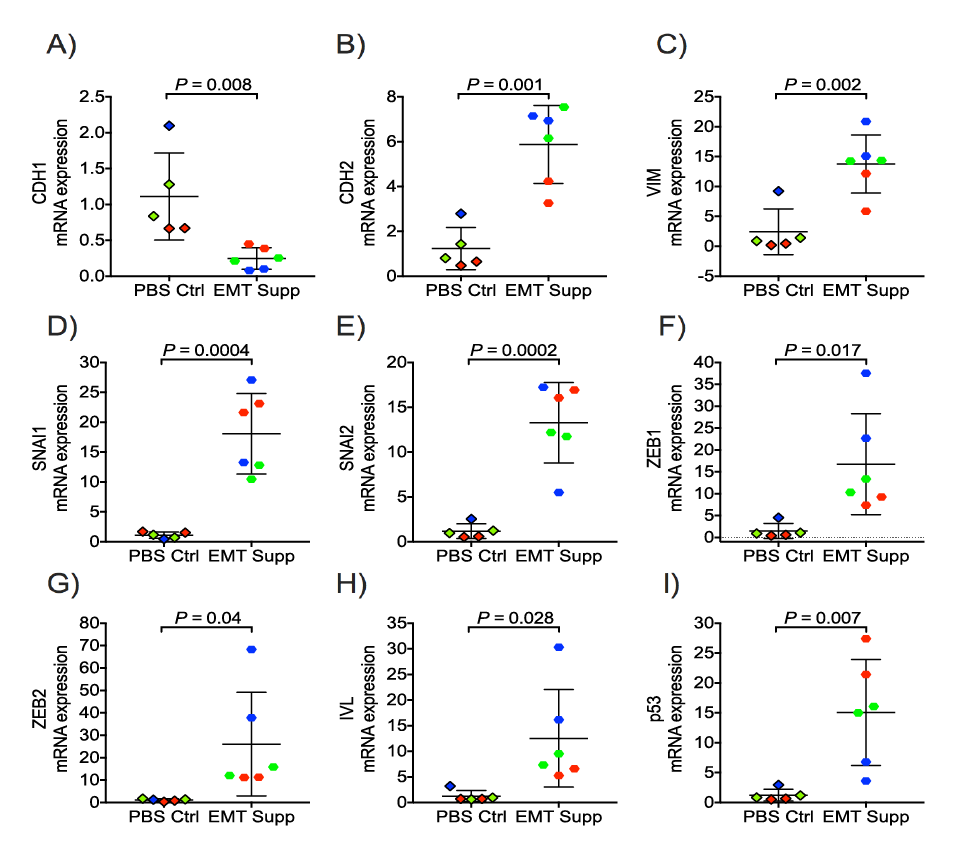
